# Short Peptides as Predictors for the Structure of Polyarginine Sequences in Disordered Proteins

**DOI:** 10.1101/2020.09.16.300582

**Authors:** B. Milorey, R. Schweitzer-Stenner, B. Andrews, H. Schwalbe, B. Urbanc

## Abstract

Intrinsically disordered proteins (IDPs) and regions (IDRs) are frequently enriched in charged amino acids. IDRs are regularly involved in important biological processes, where one or more charged residues is the driving force behind a protein-biomolecule interaction. Several lines of experimental and computational evidence suggest that polypeptides and proteins that carry high net charges have a high preference for extended conformations with average end to end distances exceeding expectations for self-avoiding random coils. Here, we show that charged arginine (R) residues in even short glycine (G) capped model peptides (GRRG and GRRRG) significantly affect the conformational propensities of each other when compared to the intrinsic propensities of a mostly unperturbed arginine in the tripeptide GRG. A conformational analysis based on experimentally determined J-coupling constants from heteronuclear NMR spectroscopy and amide I’ band profiles from polarized Raman spectroscopy reveals that nearest neighbor interactions stabilize extended β-strand conformations at the expense of polyproline II and turn conformations. The results from MD simulations with an CHARMM36m force field and TIP3P water reproduce our results only to a limited extent. The use of the Ramachandran distribution of the central residue of GRRRG in a calculation of end-to-end distances of polyarginines of different length yielded the expected power law behavior. The scaling coefficient of 0.66 suggests that such peptides would be more extended than predicted by a self-avoiding random walk. Our findings thus support in principle theoretical predictions of Mao et al. (*Proc. Natl. Acad. Sci. USA, 107, 8183-8188, 2010*).

**Significance:** Intrinsically disordered proteins are rich in charged and deficient in hydrophobic residues. High net charges of disordered protein segments favor statistical coil ensembles which are more extended than a self-avoiding random coil. It is unclear whether the chain extension solely reflects the avoidance of non-local interactions or also local nearest neighbor interactions provide significant contributions. The relevance of nearest neighbor interactions, which are neglected in random coil models, has been emphasized in the literature, but only sporadically considered in molecular modellings of disordered proteins and peptides. We determined the Ramachandran distributions of protonated arginine in GRRG and GRRRG peptides. Our results reveal the contribution of nearest neighbor interactions to the extended conformations reported for a variety of poly-arginine protein segments.

## Introduction

Intrinsically disordered proteins (IDPs) exhibit a dynamic behavior that allows them to exist as an ensemble of energetically similar albeit structurally distinct conformations under physiological conditions. Although IDPs do not have a single well-defined native structure, they are involved in many life-sustaining biological processes.^1–4^ This disconnect between structure and function has challenged the now outdated central dogma of protein biophysics that a well-defined protein structure is necessary for function. Early attempts to characterize IDPs in structural terms treated them as unfolded proteins and categorized their conformational ensembles in terms of two states, i.e. collapsed globules (in poor solvents) and self-avoiding random coils (in good solvents).^2,5–7^ NMR studies using J couplings, NOEs and dipolar couplings revealed the occurrence of local residual structure in the former that are stabilized by non-local interactions.^8–12^ The propensities of proteins for these two states depend on their amino acid composition. Since IDPs are generally rich in amino acid residues with charged or polar side chains^2,13,14^ one might suspect that water would function as a good solvent and thus favor the more extended self-avoiding coil state. However, several lines of evidence reported in the literature show that IDPs and homopeptides with polar residues adopt collapsed conformations even in the presence of denaturing cosolvents. The most prominent example is polyglutamine.^15,16^ Even an excess of charges does not guarantee an extended state. Mao et al. provided computational evidence for the notion that a preference for the coiled state requires that the net charge per residue of a polypeptide chain must exceed a certain value for moving the polymer above the theta point at a given temperature,^16^ where intramolecular and solute-solvent interactions balance each other. This notion is confirmed by single molecule experiments on protein segments and predicted scaling exponents of a large number of proteins.^14^

Conceptually both the ideal Flory random coil model and the self-avoiding random walk model are based on the assumption that a polymer can be described as a freely jointed chain of subunits.^17–20^ In the case of polypeptides/proteins this would imply a nearly unhindered rotation around the backbone torsions φ and Ψ that connect the peptide groups. However, this idealization is at variance with reality in that steric hindrance and electrostatic interactions restrict the dihedral angles φ and Ψ. This restriction is generally considered as largely insignificant for two reasons. First, the sterically allowed regions of the Ramachandran plot of individual amino acid residues are very similar with the sole exception of glycine (more extended) and proline (more restricted). Hence, individual sequences of polypeptides and proteins would not matter much with regard to experimentally accessible parameters like the radius of gyration and end-to-end distances. Second, multiple lines of experimental evidence revealed that e.g. the radii of gyration of proteins and IDPs in denaturing solvents exhibit scaling exponents which cluster around 0.59, the expectation value for a self-avoiding random walk in a good solvent.^21,22^

The above line of thinking can be questioned based on a variety of experimental and computational results. First of all, it is now well established that at the individual amino acid level, the accessible regions of the Ramachandran plot are restricted by far more factors than just steric allowance. Backbone and side chain interactions with the solvent cause the intrinsic conformational propensities of amino acids to differ significantly.^23–26^ Residues preferably sample the upper left quadrant of the Ramachandran plot (between 70 and 80%).^27–31^ The remaining 20-30% are generally distributed over several turn-like conformations. Conformational distributions differ in terms of their sampling of polyproline II (pPII) and β-strand like conformations and thus with regard to their conformational backbone entropies.^32,33^ Second, the random coil model ignores the influence of nearest neighbor residues on conformational distributions since it is based on the assumption that the conformational samplings of each amino acid in a polypeptide chain are completely independent from each other. This is the so-called isolated pair hypothesis (IPH).^18^ This hypothesis is at variance with experimental as well as bioinformatical evidence which show that conformational propensities as well as positions of basins associated with different secondary structures depend on the side chain properties of respective neighbors.^34–40^ Hence, a proper description of the underlying nearest-neighbor interactions (NNIs) has to be included in all theories of unfolded and disordered proteins.

Besides solvent-mediated effects, one possible source of NNIs could be the electrostatic interaction between charged side chains. This issue seems to be of general relevance for IDPs owing to the common occurrence of charged residues in these proteins. Since many proteins are rarely fully disordered in their native state, some of the focus in the field has shifted to defining intrinsically disordered regions (IDRs) that exist in proteins that are otherwise ordered with well-defined structures. These regions are often rich with charged amino acids that can drive interactions with other proteins or biomolecules. The widespread role of IDRs in biomolecular interactions suggests they are easily recognizable and accessible to their binding partners. Patches of charged amino acids often exist as so-called *linear motifs*, which is a class of IDRs.^41–44^ As indicated above, protein segments and polypeptides with a high net charge per residue can be more extended than predicted by a self-avoiding random coil model.^16^

The study described in this paper was motivated by a recent investigation of 20 protamine sequences which are all very rich in arginine.^16,19^ The sequences differed with regard to their net charge and the distribution of arginine residues. Atomistic simulations of these polypeptides strongly suggest that an increase in net charge above a certain threshold value can trigger a globule to coil transition. Scaling coefficients for peptides with very high net charges were predicted to exceed the canonical 0.59 value, suggesting that the classical random coil model does not suffice in these cases. Mao et al. described the extended backbone of the investigated poly-arginine as a rod-like structure.^16^ In their simulation these authors used a molecular mechanics force field based on OPLS-AA/L parameters and an implicit solvent model. In order to account for electrostatic interactions between side chains a mean field electrostatic energy term was added to the total energy function.

In this study, we investigate whether the predicted non-random coil structure of polyarginine could be related at least in part to NNIs between arginine residues. A previously reported conformational analysis of arginine in the cationic model peptide Gly-Arg-Gly (GRG) showed that this amino acid has a comparatively high propensity for the pPII structures with a smaller but still significant tendency to populate more extended β-strand-like conformations.^45^ In line with findings for other GxG peptides with polar or charged guest residues x the population of turn-like conformations is not negligible (around 21%).^28,29^ Recent density functional theory (DFT) calculations have provided evidence that the hydration shell around the peptide backbone of GRG stabilizes this conformational bias.^46^ Here we combine data from experimental NMR and polarized Raman spectroscopic experiments to perform a conformational analysis on individual arginine residues in GRRG and GRRRG to probe the effect that neighboring charged arginine residues have on each other. Our experimentally based results are compared to distributions produced by MD simulations with an CHARMM36m force field combined with a TIP3P water model.^47^ We relate our findings to the predictions of Mao et al.^16^ by using them to predict the scaling law for a poly-arginine chain of 25 residues.

## Material and Methods

### Solution NMR experiments

For ^1^H NMR experiments, peptides were purchased from Genscript. They were dissolved in an aqueous solution of 90% H_2_O / 10% D_2_O at a concentration of 100 mM, and the pH was adjusted to < 2.0. The measurements were recorded with a 600 MHz Bruker AV600 spectrometer. For all other NMR experiments, isotopically labeled amino acids were purchased from Cambridge Isotopes, and the peptides of interest were synthesized using an Applied Biosystems 433A peptide synthesizer. Purification was performed via reverse-phase HPLC before freeze drying. The product of the solid phase peptide synthesis was checked using electrospray ionization (ESI) and matrix-assisted laser desorption/ionization (MALDI) mass spectrometry. The labeled peptides were dissolved at a concentration of between 5 to 10 mM in an aqueous solution of 90% H_2_O / 10% D_2_O, and the pH was adjusted to < 2.0. The measurements were recorded with a Bruker 800 MHz AV800 spectrometer. Table S1 lists the measured J-coupling constants with the homo- and heteronuclear NMR technique that was utilized as well as the technique employed for a specific J-coupling constant. A detailed description of the application of each individual technique to short model peptides are given in an earlier papers.^1,2^

### Raman experiments

Peptides were dissolved in D_2_O to either 100 (GRRG) or 70 (GRRRG) mM concentration. The pH was adjusted to <2.0 with deuterium chloride (DCl). The Raman spectra were obtained with the 514.5 nm radiation of a Spectra-Physics (Mt. View, CA, USA) Argon laser (200 mW). The laser beam was directed into a Renishaw confocal microscope and focused onto a thin glass cover slip with a 20x objective. The scattered light was filtered with a 514.5 notch filter. Polarized Raman spectra were recorded using a Renishaw confocal microscope. The microscope is equipped with a linear polarizer and a λ/2 plate which rotates the y-polarized light into the x-direction. Polarized and unpolarized spectra were recorded between 1400 and 1800 cm^-1^. A background spectrum of the D_2_O/DCl mixture was recorded on the same day as and later subtracted from each peptide spectrum. The unpolarized spectra were recorded as an average of five measurements using the WiRE 3.3 Renishaw software, and the polarized spectra were an average of ten total measurements.

### J-coupling constants and Karplus parameters

This section describes the mathematical basis for our conformational analysis. The Karplus equations relate certain J-coupling constants to the dihedral angles of the peptide backbone, φ and Ψ. These equations take the form of eq. 1,

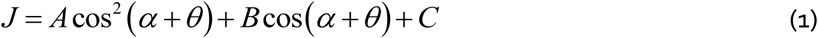

where *J* is the J coupling constant, A, B and C denote empirical Karplus parameters, α=φ,Ψ represents the peptide backbone dihedral angle to which the respective J-coupling constant is related, and *θ* is a phase correction factor. Though the Karplus equations all take the same form, differences in Karplus parameters and phases cause each J-coupling constant to depend on the backbone angle in a different manner. See Figure S1 for a visual representation of the Karplus curves.

Different sets of Karplus parameters have been reported in the literature. As in our earlier studies we used the empirical values of Wang and Bax for φ-dependent coupling constants.^3^ Other parameter sets reported by the Bax-group yield very similar Karplus curves.^4,5^ For reasons detailed below we also used Karplus parameters which were obtained with DFT calculations for alanine-based peptides for ^3^J(H^α^,C’).^6^ For ^1^J(N,C^α^), we use the parameters reported by Wirmer and Schwalbe.^7^ In view of the importance of this coupling constant for an assessment of conformational distribution along the Ψ-axis we additionally performed fits with the parameters reported by Ding and Gronenborn.^8^ The ^2^J(NC^α^) is peculiar in that it depends on the Ψ-angle of the preceding residue of the amide nitrogen. Table S2 lists the Karplus parameters used for our analysis.

*

### Raman band profiles

In what follows we will use the term amide I if we characterize this peptide mode in general terms. Amide I’ refers to the band in spectra of peptides dissolved in D_2_O. By substituting H_2_O by D_2_O we eliminate the NH- and HOH bending contribution to the eigenvector of amide I.^9,10^ The excitonic coupling model used to simulate the amide I’ profiles in the polarized Raman spectra of the investigated peptides have been described in several of our earlier publications.^11,12^ Briefly, the single excitation excitonic states of amide I are described as linear combinations of local eigenstates of local harmonic wavefunctions. The eigenenergies (wavenumbers of amide I bands) and the coefficients of the above linear combination are obtained from the diagonalization of the Hamiltonian which for a pentapeptide is written in the basis of the excited eigenstates of the respective local amide I vibrations as follows:

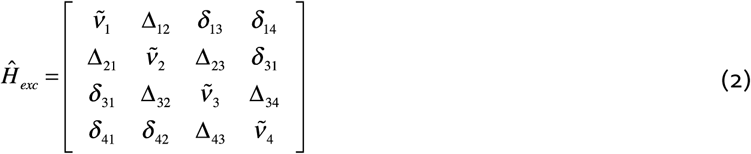

The corresponding Hamiltonian of a tetrapeptide is a 3 × 3 matrix. The diagonal elements 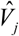 are the vibrational frequencies of the local amide I oscillators. Δ_*ii*±1_denotes nearest neighbor coupling energies that are φ and Ψ dependent. The algorithm used to calculate Δ_*ii*±1_is described by Schweitzer-Stenner.^13^ Non-nearest coupling *δ*_*ij*_ can be reasonably well described by transition dipole coupling.^14,15^ Once the Schrödinger equation is solved, the obtained eigenfunctions can be used to calculate the Raman tensor of the excitonic states. To this end Raman tensors of the local oscillators have to be transformed into one common coordinate system which is centered at the C-terminal nitrogen of the peptide.^12^ This brings about a conformational dependence of the Raman tensor. In this coordinate system the Raman tensor is a 3×3 matrix from which the intensities of polarized Raman scattering for the *m*^*th*^ excitonic transition can be obtained as follows:

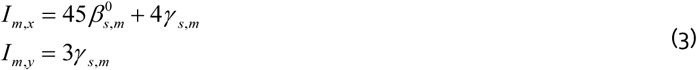

The subscripts x and y indicate polarization with regard to polarization of the exciting laser beam. The tensor invariants 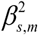 and 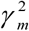 are calculated as:

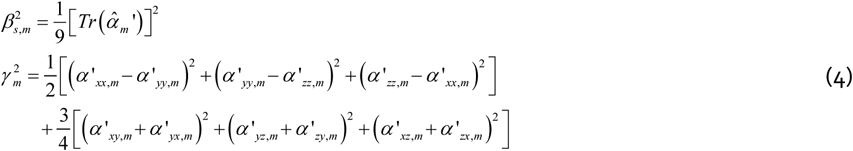

where *α* ‘_*kl,m*_ (*k,l=x,y,z*) denote the elements of the Raman tensor. The prime sign indicated that each tensor element is related to the derivate of the molecular polarizability with respect to the normal mode of amide I. The tensor invariants reflect the average over all molecular orientations in the laboratory coordinate system.

### Conformational analysis part I: J-coupling

Ramachandran distributions of amino acid residues are expressed in terms of linear combinations of weighted two-dimensional Gaussian sub-distributions associated with known secondary structures:

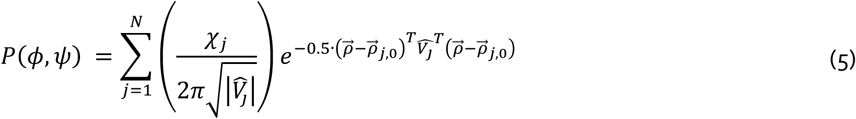

where the index *j* represents the *j*^*th*^ conformation summed over N different sub-distributions with their own statistical weights χ_j._ The position vector 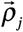 represents a coordinate pair in the Ramachandran plot, while 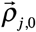 denotes the coordinates of the maximum of the *j*^*th*^ distribution. The matrix 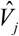 contains the standard deviations with regard to φ and Ψ as on-diagonal and correlation coefficients as off-diagonal elements. The latter have been assumed to be zero for the analyses described in this paper.

The J-coupling constants were calculated as the ensemble average over the above distribution:^13^

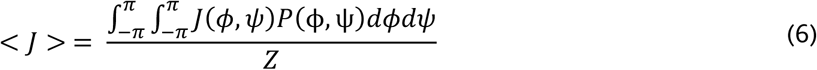

where *o* represents the considered spectroscopic observable *J, I*_*x*_ or *I*_*y*_. Z is the partition sum.

In order to obtain a quantitative criterion for the comparison of Ramachandran distribution we utilize the Hellinger distance:^16,17^

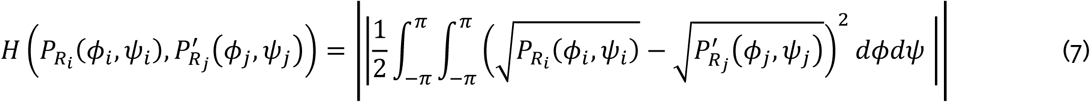

The Hellinger distance measures the dissimilarity of two two-dimensional distributions. If the value is zero, the compared distributions are identical. A value of 1 indicates orthogonality. However, since individual residues sample a restricted space of the Ramachandran plot H will never reach high values for statistical coil conformations. Here, we adopt the criteria of Schweitzer-Stenner and Toal^17^ according to which two distributions are very similar if *H* ≤ 0.1. Values with 0.1 < H ≤ 0.25 and 0.25 < H ≤ 0.4 as moderately similar and dissimilar, respectively. Any larger value indicates significant dissimilarity.

In addition to our use of the Hellinger distance we compared the Ramachandran distributions of the GRG and the investigated peptides by calculating their conformational entropy. For individual residues it is calculated as:

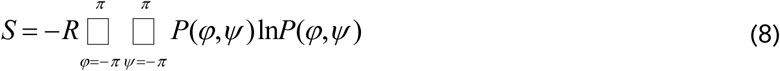

where R is the gas constant. In the absence of any nearest neighbor interactions the total entropy of a peptide would be the sum of individual entropies of the respective residues. If, as expected, nearest neighbor interactions affect the conformational distributions of GRRG and GRRRG, the total conformational entropy of a peptide would reflect correlation effects which can be accounted for by conditional probabilities.^18–20^ The calculation of such entropies is outside of the scope of this paper.

### Conformational analysis part II: Amide I

Since the J-coupling constants are site-specific with regard to their conformational dependence, they can be individually calculated by using the Ramachandran plot for each individual residue. Since the structure sensitivity of amide I’ results from coupling between oscillators in different residues this is not an option for the simulation of amide I’ band profile. Since a full consideration of the combined Ramachandran space of two and more residues would be computational demanding we used a truncated distribution function for which we focus sampling on the areas of the Ramachandran plot for which we expect significant conformational sampling. A total of 11 conformers represents the pPII and β-strand troughs. Each are located within a quadrant around the sub-distribution maxima obtained from the primary analysis of J-coupling constants and carries 1/11 of the total statistical weight of the conformation. Hence, we substitute the two-dimensional Gaussian distributions by cuboids with widths of 2*Δϕ_j_ and 2*ΔΨ_j_ in φ and Ψ direction, where Δϕ_j_ and ΔΨ_j_ were set equal to the standard deviations of the respective Gaussian distributions. The height of the cuboids is given by the statistical weights of the conformations divided by the number of representing data points. Hence, the probability distribution reads as:

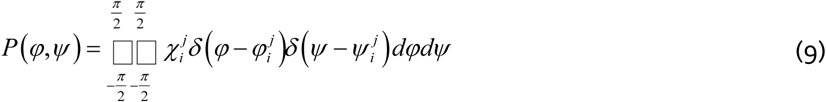

where 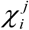 is the mole fraction of the *i*^*th*^ data point of the *j*^*th*^ sub-distribution, which exhibits the Ramachandran coordinates 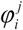 and 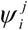 and δ indicates the Dirac delta function. The amide I’ band profiles of this statistical ensemble are then calculated as:

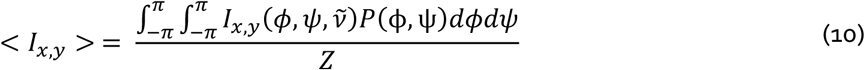

where

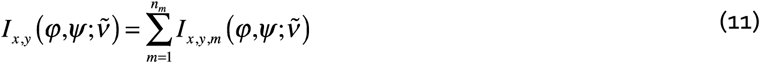

and 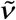 denotes the wavenumber position in the spectrum.

### Calculating the average peptide length

We used the distance between the N- and C-terminal carbonyl oxygen as a measure of the peptide’s length. We calculated it as follows. The vector connecting the CO group of two adjacent residues can be obtained by the vector addition

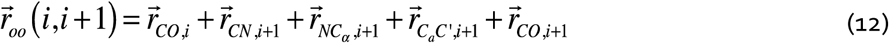

where the bond with which a vector is associated is indicated as subscript. Apparently, the contribution of the last two vectors in the expression to the total distance depends on the two dihedral angles of the (*i+1)*^*th*^ residue. Each vector 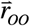 is transformed from the respective coordinate system of its residue into a common coordinate system situated at the C-terminal residue. After the transformation, all vectors can be added up to yield the distance between the N -terminal and C-terminal residues.

### Calculation of Debye length

In order to gauge the contribution of electrostatic interaction between charged groups (arginine side chains and the N-terminal in the present case) could be of any significance for an understanding the results of our conformational analysis we have to estimate the Debye length for the solution conditions used in our study. To this end we employ

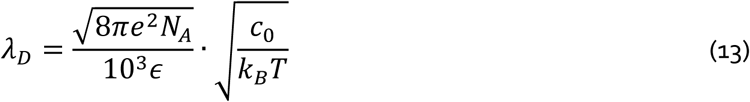

where N_A_ is the Avogardo number, *e* is the elementary charge, *k*_*B*_ is the Boltzmann constant, T the absolute temperature and *c*_*0*_ the molar ion concentration which is very much determined by the peptide concentration.

### MD simulations and mesostate calculations

Molecular dynamics (MD) simulations were performed using GROMACS 5.1.2.^21–23^ on single GRG, GRRG and GRRRG peptides with an CHARMM36m force field and an TIP3P water model.^24^ Additional simulations using Amber ff14SB and OPLS-AA/M with TIP3P and TIP4P, respectively, were carried out for GRG. For the simulation with CHARMM36m, the peptides are capped with protonated NH_3_ ^+^ termini and neutral C-termini (COOH) to mimic the influence of the acidic conditions used for the NMR and Raman experiments. For each simulation, a single peptide was solvated in a cubic box of 40°40°40 Å^3^. The energy minimization is performed using steepest descent minimization for 100,000 steps, followed by a pressure equilibrium for 20 ps at 300 K and a pressure of 1 bar. Stoichiometric amounts of Cl^-^ atoms are added to ensure electroneutrality. Each 500 ns long trajectory is acquired using the velocity rescale thermostat^25^ and the Berendsen barostat.^26^

In order to assess the MD-based Ramachandran distributions we calculated the occupation of mesostates that cover populated regions of the Ramachandran plot. They are visualized in Figure S2. They are defined as follows: (a) pPII(−90° < ϕ < -42°, 100° < Ψ < 180°), (b) anti-parallel β-strand (aβ) (−180° < ϕ < -130°, 130° < Ψ < 180°), (c) parallel β-strand (pβ) (−150° < ϕ < -130°, 100° < Ψ < 128°), (d) the transition region between pPII and aβ(βt) (−130° < ϕ < -90°, 130° < Ψ < 180°), and (e) right handed helical (−90° < ϕ < -32°, -60° < Ψ < 14°). The mesostate populations were calculated from the MD trajectories using time frames 50-500 ns as the number of conformations within each mesostate region normalized by the total number of conformations. Eq. 6 was used to calculate the J-coupling constants for each of non-terminal residues of the investigated peptides.

## RESULTS AND DISCUSSION

Our conformational analysis of the non-terminal residues of GRRG and GRRRG is based on a previously described model which describes the Ramachandran plot of individual residues as a superposition of two-dimensional Gaussian distributions associated with known secondary structures. The thus obtained probability density distribution is optimized to reproduce experimentally obtained J-coupling constants and amide I’ Raman profiles. Details are described in the Material and Methods section.

### Gaussian modelling of GRRG and GRRRG

The construction of our Ramachandran plots started with six experimentally determined J-coupling constants listed in Table S1. They were selected because of their rather different dependencies on the dihedral angles φ and Ψ of the peptide backbone. Each coupling constant can be related to one of these dihedral angles with a Karplus equation.^49,50^ These equations all take the same form as eq. 1, but the differences in phase and Karplus parameters cause each J-coupling constant to depend on the backbone angle in a vastly different manner. Figure 1 shows the NMR spectra measured to determine the indicated coupling constants for GRRG. The corresponding experimental values obtained from this and the corresponding spectra of GRRRG are listed in Table 1. The measured coupling constants represent conformational averages as described in the Material and Methods section of the Supporting Information. We optimized the model by minimizing the difference between experimental and simulated J-coupling constants, which reflect the conformational average of the two dihedral angles of the peptide backbone (φ and Ψ). The purpose of the fitting is to produce population statistics for five possible peptide conformations, which can be displayed as distinct regions on a Ramachandran plot. Each of these conformations is represented by a two-dimensional Gaussian distribution. Based on earlier results obtained for fully protonated GRG^45^, we selected the following conformations: (a) pPII with -85°<ϕ<-65° and 140°<Ψ<180°, (b) β-strand with -100°<ϕ<-180° and 140°<Ψ<180°, (c) right-handed helical with -80°<ϕ<-50° and -40°<Ψ<0°, (d) inverse γ-turn with -90°<ϕ<-70° and 70°<Ψ<80°, (e) left-handed helical with 50°<ϕ<80° and 60°<Ψ<90°, and (f) the asx-turn conformation with 60°<ϕ<-90° and 130°<Ψ<180°. The latter is not populated in the Ramachandran plot of GRG (Figure 2a), but occurs frequently in Ramachandran plots of residues with hydrogen bonding capability.^28,29^ The rather large β-strand region encompasses all three β-strand mesostates introduced above. We neglected the γ-turn conformation in the lower right quadrant of the Ramachandran plot because of its low population in GRG and the necessity to limit the number of free parameters.

**Table 1.** Experimental and computed J-coupling constants of the indicated arginine residues of GRG, GRRG and GRRRG in aqueous solution. Experimental uncertainties are listed in parenthesis. The theoretical values were calculated with a Gaussian distribution model described in the text.^48^ The mole fraction of the considered conformations are listed at the bottom of each column.

|  |  | GRG* | GRRG |  | GRRRG |  |  |
| --- | --- | --- | --- | --- | --- | --- | --- |
|  |  | *Ri* | *R1* | *R2* | *R1* | *R2* | *R3* |
| $^3J(H^N H^\alpha)$ | *exp.* | 6.66<br>(0.01) | 6.54 (0.017) | 7.22 (0.035) | 6.55 (0.040) | 6.65 (0.044) | 7.14 (0.052) |
|  | *Gaussian* | 6.47 | 6.51 | 6.92 | 6.66 | 6.65 | 7.00 |
|  | *MD* | 6.74 | 6.28 | 6.8 | 6.2 | 6.53 | 7.45 |
| $^3J(H^N C')$ | *exp.* | 1.00<br>(0.08) | 1.01 (0.067) | 1.26 (0.151) | 1.01 (0.067) | 1.41 (0.403) | 1.08 (0.201) |
|  | *Gaussian* | 0.98 | 0.96 | 1.21 | 1.09 | 1.60 | 1.17 |
|  | *MD* | 1.11 | 1.26 | 1.08 | 1.25 | 0.97 | 0.94 |
| $^3J(H^\alpha C')$ | *exp.* | 2.47<br>(0.05) | 2.02 (0.067) | 0.806<br>(0.201) | 1.61 (0.269) | 0.672<br>(0.067) | 0.806 (0.067) |
|  | *Gaussian* | 2.36 | 1.87 | 1.79 | 1.72 | 1.7 | 1.85 |
|  | *MD* | 2.15 | 2.43 | 2.34 | 2.39 | 2.7 | 2.39 |
| $^3J(H^N C^\beta)$ | *exp.* | 1.79<br>(0.15) | 1.87 (0.220) | 1.49 (0.660) | 1.32 (0.220) | 0.99 (0.330) | 1.10 (0.440) |
|  | *Gaussian* | 1.75 | 1.81 | 1.44 | 1.64 | 1.32 | 1.43 |
|  | *MD* | 1.61 | 1.69 | 1.6 | 1.73 | 1.79 | 1.4 |
| $^1J(NC^\alpha)$ | *exp.* | 11.02<br>(0.10) | 11.79<br>(0.214) | 11.60<br>(0.165) | 11.92<br>(0.139) | 11.82<br>(0.146) | 11.59 (0.142) |
|  | *Gaussian* | 10.94 | 11.57 | 11.46 | 11.6 | 11.53 | 11.36 |
|  | *MD* | 11.16 | 11.29 | 11.19 | 11.34 | 11.03 | 11.09 |
| $\chi_{pII}$ | | 0.58 | 0.62 | 0.46 | 0.57 | 0.34 | 0.40 |
| $\chi_\beta$ | | 0.20 | 0.17 | 0.34 | 0.29 | 0.46 | 0.39 |
| **Total Extended** |  | **0.78** | **0.79** | **0.80** | **0.86** | **0.80** | **0.79** |
| $\chi_{rhHel}$ | | 0.08 | 0.08 | 0.04 | 0.14 | 0.19 | 0.21 |
| $\chi_{iy}$ | | 0.03 | 0.06 | 0.15 | 0.00 | 0.00 | 0.00 |
| $\chi_{lhelix}$ | | 0.08 | 0.00 | 0.00 | 0.00 | 0.00 | 0.00 |

\*The probability distribution of *R1* in GRRG contains also a 6% asx-turn population.

**Figure 1:**
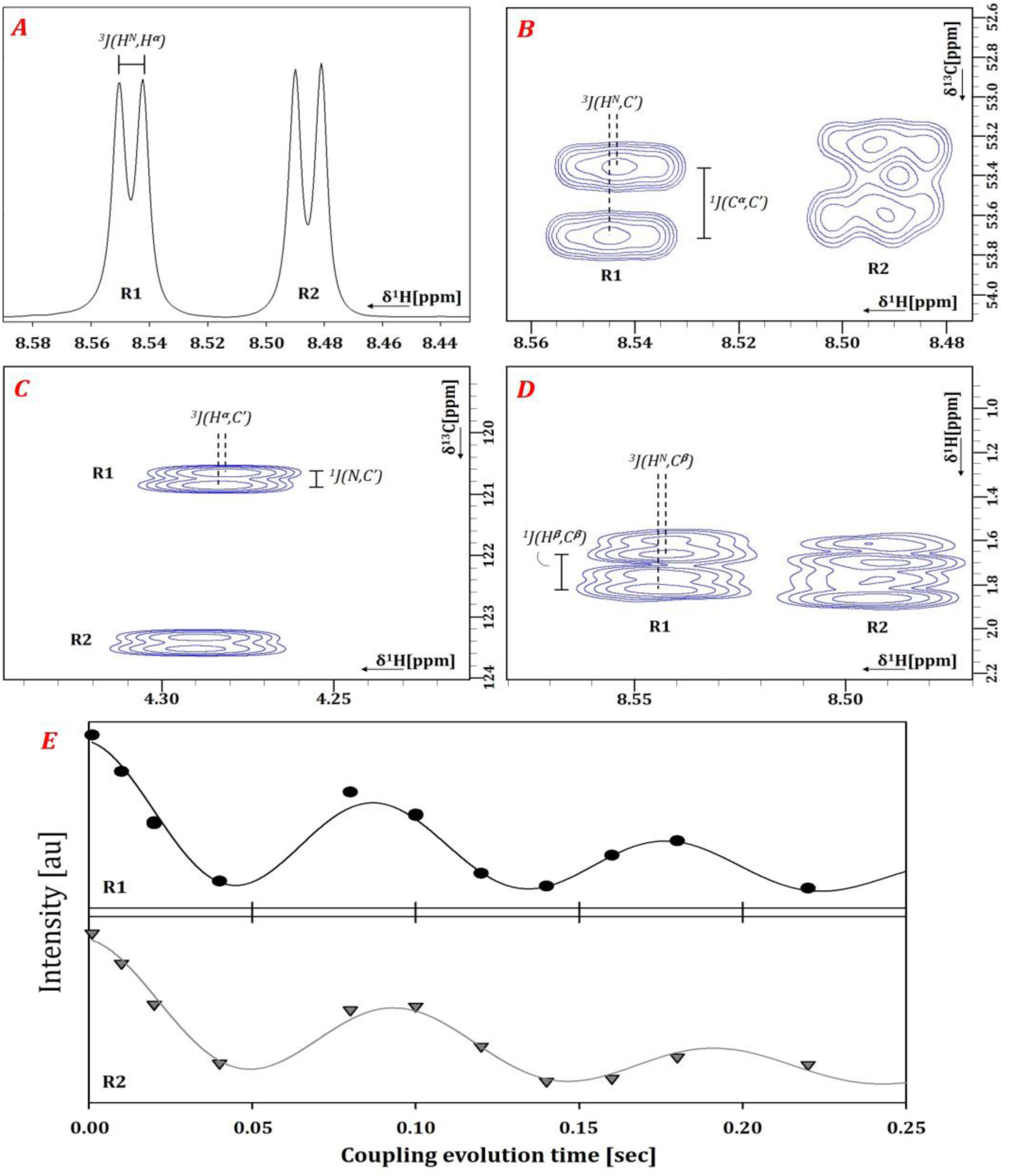
NMR measurements to determine J-coupling constants, shown for GRRG. A) Experimental doublets resulting from the ^3^J(H^N^,H^α^) coupling. B) Soft HNCA-COSY for determination of ^3^J(H^N^,C’). C) CO-coupled [H]NC_α_H_α_ for the determination of ^3^J(H^α^,C’). D) HNHB[HB]-E.COSY for the measurement of ^3^J(H^N^,C^β^). E) Intensities obtained from a J-modulated ^1^H,^15^N-HSQC experiment and the fit *I* (*τ*) = *A*cos(^1^ *J* (*N,C* ‘)*τ*)cos(^2^ *J* (*NC* ‘)*τ*)×exp(−*τ T*_*2*_) to determine ^1^J(N_i_,C^α^_i_) or ^2^J(N_i_,C^α^_i-1_) are shown in gray.

**Figure 2.**
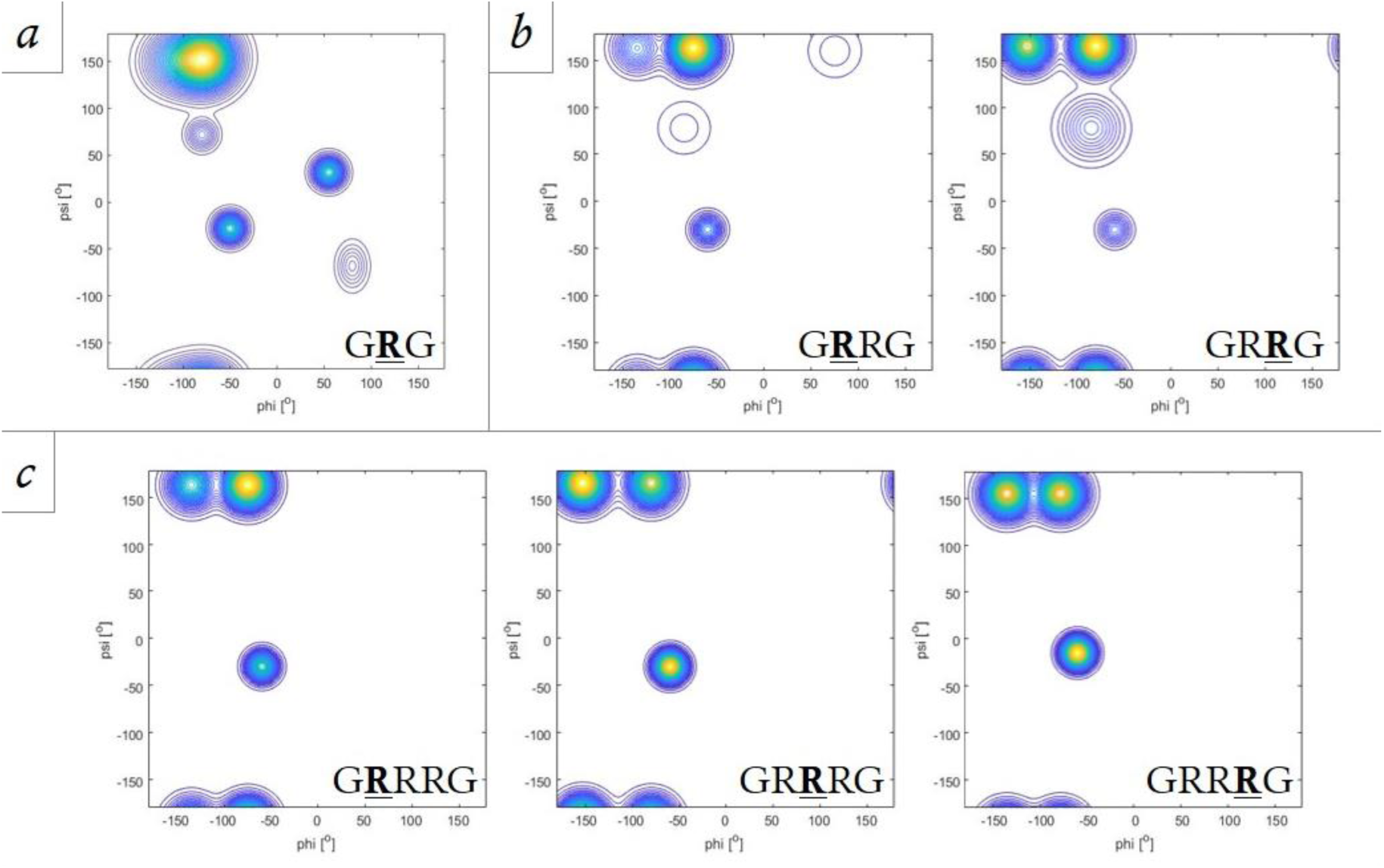
Ramachandran plots for individual arginine residues in GRG, GRRG and GRRRG derived from a conformational analysis of these peptides as described in the text. The bold capital R indicates the residue for which the conformational distribution is displayed.

To extract the relative populations of each of the considered conformations, a set of two-dimensional Gaussian functions were centered around (φ,Ψ) coordinates in the above regions of the Ramachandran space. The relative populations were used as free parameters subject to normalization in a non-linear least square fit using the *lsquarefit* module of Matlab 2019b. Initial φ- and Ψ-positions and halfwidths were taken from the Gaussian model used for GRG. The respective position values of the three major sub-distributions (based on the obtained mole fractions) were varied to obtain the best fit to our experimental J-coupling constants by using individual Karplus equations, the Karplus constants (Table S2) and the formalism described in Material and Methods. The fitting was carried out for each of the individual arginine residues. In a first round we used the Karplus parameters reported by Wirmer and Schwalbe for ^1^J(N,C^α^).^51^ In a second round we used the parameter set for ^1^J(N,C^α^) reported by Ding and Gronenborn.^52^ Checking the performance of these different parameter sets was deemed important because the ^1^J(N,C_α_) values of GRRG and GRRRG were found to be systematically larger than that of GRG, thus indicating that NNIs cause an upshift of the distributions and/or a redistribution of mole fractions between turn-like and extended conformations. Assessing these changes correctly is of utmost importance of the specific aims of this study.

To check the validity of the obtained Gaussian parameter values we used them to calculate the x- and y-polarized Raman profile of the respective amide I’ mode. To this end we used the truncated distribution model described in Material and Methods. We always calculated the J-coupling constants for the truncated distributions and found them always practically identical with the ones obtained with the complete probability distributions.

The experimentally determined and simulated amide I’ band Raman profiles for GRRG and GRRRG are shown in Figure 3. In each case, the agreement between experimental and computed profiles is satisfactory. Slight differences between e.g. the experimental perpendicular polarized Raman scattering and the calculated profile might reflect minor calibration issues.

**Figure 3.**
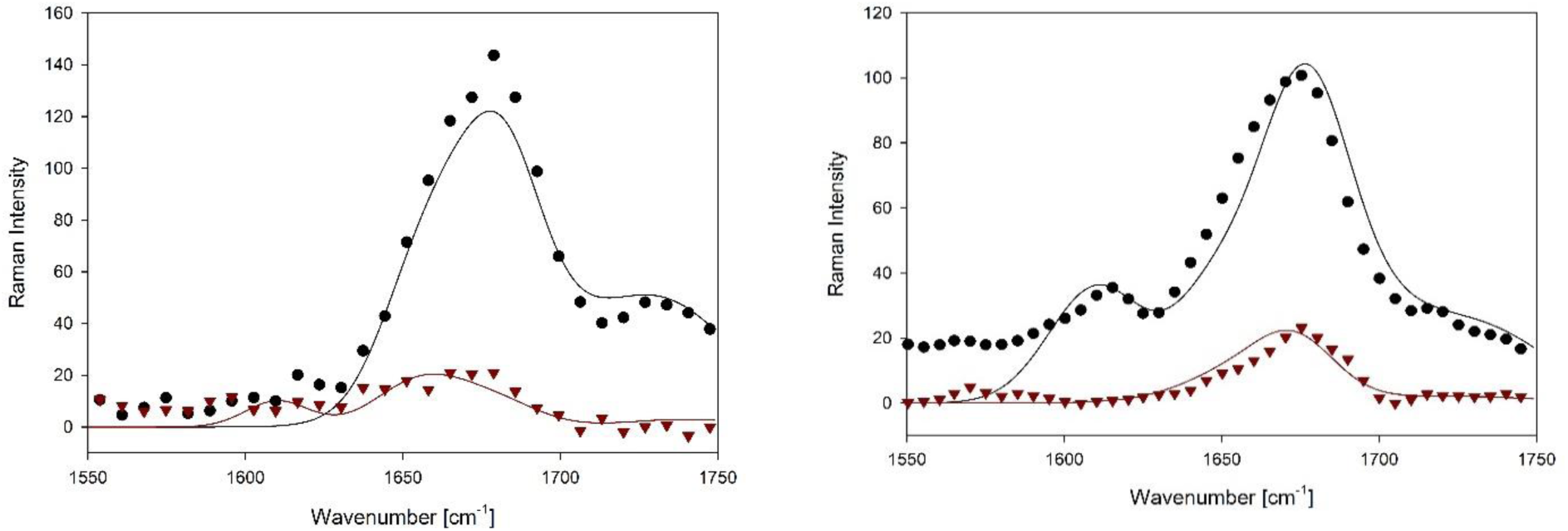
Amide I’ profiles in the x-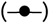 and y-polarized 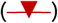 Raman spectra of GRRG and GRRRG in D_2_O at pD 2. The solid lines in the figures result from simulations based on conformational distributions inferred from J-coupling constants.

In earlier studies we had added the amide I’ in IR and vibrational circular dichroism (VCD) to our arsenal of experimental data. VCD profiles were shown to be particularly sensitive to changes in Ψ in the pPII region. Unfortunately, we had to refrain from using these tools for the current study. Gosh et al. showed that the symmetric and asymmetric CNC stretching modes couple through space with the two amide I’ modes of the arginine dipeptide.^53^ Since the N-terminal amide I’ modes of GRG is significantly blueshifted relative to its position in the arginine dipeptide spectrum, it is moved out of the region where excitonic coupling with the arginine modes matter. However, the C-terminal mode is still in the region where coupling can occur. IR and VCD spectra taken in our laboratory show that this happens to some extent for GRRG and GRRRG, but for some reason not in GRG. Such coupling is unlikely to significantly affect the Raman profiles, though, since these arginine modes are not Raman active. Therefore, we confined ourselves to the use of polarized Raman profiles in the current study.

The J-coupling parameters and the respective reduced χ_J_^2^ values are listed in Table 1. The latter were calculated as described by Zhang et al.^24^ It should be noted that ^2^J(N,C^α^) was not incorporated in the fitting process. The reported values were calculated using the Ramachandran distributions obtained from the fit to the other coupling parameters. One might be inclined to argue that the χ_J_^2^ values obtained for R2 of GRRG and for R2 and R3 of GRRRG are too large to deem the respective fitting satisfactory. However, a closer look at the origin of these numbers is required for an appropriate assessment. A substantial contribution to these large numbers results from the difficulty in reproducing the surprisingly low ^3^J(H^α^,C’) values for the above arginine residues. The second largest contribution stems from differences between experimental and computed ^3^J(H^N^,C^β^) values, which are also comparatively low for R2 and R3. All these values are substantially different from those reported for GRG and also from the respective values obtained for the R1 residues of the two investigated peptides. The Karplus curves of these parameters in Figure S1 suggest that the low values for ^3^J(H^α^,C’) are due to conformational sampling around φ = -50° and/or -180°. The former would be to some extent compatible with the rather well reproduced ^3^J(H^N^,H^α^) and ^3^J(H^N^,C’) values, but at variance with the values observed for ^3^J(H^N^,C^β^) which exhibits a maximum in this region. A φ-value of -180° is problematic because a major population in this region would totally jeopardize the calculation of ^3^J(H^N^,H^α^) and ^3^J(H^N^,C’). For ^3^J(H^α^C’) another difficulty arises from the fact that the lowest possible value (1) is larger than our experimental values. Such low experimental value are actually not unusual for proteins as documented by Hu and Bax.^54^ All these observations strongly suggest that the reported empirical parametrization of the Karplus curve for ^3^J(H^α^,C’) becomes inaccurate at its minimum. We obtained a slightly better result with the Karplus parameters obtained from DFT calculations for alanine peptides which are listed in Table S2^55^. It is more likely that the real Karplus curves become negative in this region. This is a result that the respective empirical Karplus scenarios do not account for.^24^ It is worth mentioning in this context that a similar problem arose with earlier attempts to reproduce the low values of the ^3^J(C’,C’) coupling constant of trialanine.^49,56^ Generally, it should be noted that the utilized Karplus parameters are averages over residues with rather different side chains. At present, we do not know to what extent Karplus parameters constant for specific residues deviate from their respective average value. This side chain dependence can be expected to be significant for ^3^J(H^N^,C^β^) since it directly involves a side chain atom.

The agreement between calculated and measured values for ^3^J(H^N^,H^α^), ^3^J(H^N^,C’) and ^1^J(N,C^α^) is very satisfactory. It deserves to be noted that the Karplus curves of the two ^3^J coupling constants are nearly out of phase,^24,49^ hence our capability to reproduce both values well is of substantial significance. The differences between coupling parameters obtained with fits using the Wirmer-Schwalbe^51^ and Ding-Gronenborn^52^ parameters for ^1^J(N,C^α^) are insignificant if one considers the fits to all five R-residues in the investigated peptides. In what follows we focus on the results obtained with the Wirmer-Schwalbe parameters. Finally, the calculated values for ^2^J(N,C^α^) are systematically lower than the corresponding experimental values. This is not surprising since the corresponding Karplus plots never exceed 8.7 Hz.^49^ It deserves to be notified that the Karplus parameters for ^2^J(N,C^α^) might exhibit some significant side chain dependence as well. However, the very fact that some of the measured values are even larger strongly corroborate the predominance of extended structures. Hence, the calculated values are as close to the experimental values as possible with the used Karplus parameters.

The Ramachandran plots of GRRG and GRRRG as well as the earlier reported plot for GRG^45^ are shown in Figure 2. It is reasonable to assume that the terminal glycine residues produce minimal NNIs to the conformational ensemble of arginine, so we can use it as a good model of the intrinsic propensities of arginine. We will therefor use “Ri” when referring to the “intrinsic arginine.” The Ramachandran plot nicely illustrates the previously reported strong intrinsic preference for pPII.^45^ While the pPII basin is the most occupied one in the upper-left region of the Ramachandran plot, the broad population density actually reflects an overlap between pPII and β_t_. Ri also exhibits a nearly equal propensity for right- and left-handed helical conformations and weak though not negligible population of γ-turn conformations. The mole fractions of the respective conformations are listed in Table 1. This Ramachandran plot, the corresponding Gaussian sub-distributions, and their propensity values will be used as the basis to compare the Ramachandran plots of arginine residues in the model peptides GRRG and GRRRG (figures 1b and 1c). The comparison will reveal to what extent the propensities observed for Ri are altered due to the presence of neighboring arginines.

We first assess the differences between the conformational distributions of each arginine by looking closely at the way the populations change. First, it should be noted that pPII propensities of the N-terminal arginines, R1, in both GRRG and GRRRG (0.62 and 0.57, respectively) are just about the same as for Ri (0.58). Furthermore, all arginines exhibit a similar total propensity for extended structures pPII and the β-strand distribution which is located at the right boundary of the aβ mesostate. The total population of extended conformations falls between 0.79 and 0.86, as shown in Table 1. Irrespective of these similarities, the differences between the Ramachandran plots of Ri and R1 are noteworthy for both GRRG and GRRRG. The β-strand conformation has shifted to the left towards more negative φ-values. pPII and β-strand have both shifted up to larger Ψ-vales, as reflected by the larger ^1^J(NC^α^) values. Moreover, the population of the left-handed helical structures in GRG has been replaced by some minor sampling of asx-turns in the distribution of R1 in GRRG.

Differences between Ri and R2/R3 (GRRRG) seem to be more pronounced. A fraction of the population is redistributed from pPII to β-strand in the latter. This effect is most pronounced for R2 of GRRRG where the β-strand becomes the dominant basin. It is clearly shifted to more negative φ-values. This change is again particularly pronounced for R2, where the β-strand maximum is now at -153°, which positions it in the aβ-region. R2 and R3 do not show any population in the right half of the Ramachandran plot. Instead, the helical fraction has actually increased from the apparently significant NNI-induced redistribution.

In addition to the above comparisons of statistical weights and positions of local maxima of the probability density distribution we evaluated the dissimilarity between the conformational distributions of the doublet and triplet residues with the intrinsic propensities of Ri using the Hellinger distance (see eq. 6 in the Methods section). This metric has been used previously to quantitatively compare two Ramachandran plots.^37^ Our calculated Hellinger distances are all summarized in Table 2. For both the twin and triplet arginine residues the Hellinger distance (for comparisons with Ri) ranges from 0.33 to 0.35, indicating that all arginines are moderately different from Ri (cf. the criteria in Material and Methods). A comparison of the doublet arginines results in a Hellinger distance of 0.13, suggesting that they are much more similar to each other than they are to Ri. For the arginine triplet we obtained the following Hellinger distances: 0.13 for R1/R2, 0.13 for R1/R3, and 0.15 for R2/R3. All the values suggest that the compared distributions are moderately similar. These values reveal again that the changes produced by a single neighbor are more pronounced than the one induced by the presence of a second interacting neighbor. However, in spite of the rather low Hellinger distances for R1/R3 and R2/R3 the increased β-strand propensity of R2 is noteworthy and deserves further consideration (*vide supra*).

**Table 2.** List of Hellinger distances comparing the Ramachandran plots of the indicated residues in the investigated peptides.

|  |  | Ri |  |  |
| --- | --- | --- | --- | --- |
| GRRG | R1 | 0.34 | R1 |  |
|  | R2 | 0.35 | 0.13 |  |
|  |  | Ri |  |  |
| GRRRG | R1 | 0.34 | R1 |  |
|  | R2 | 0.35 | 0.13 | R2 |
|  | R3 | 0.33 | 0.13 | 0.11 |

Another parameter that can be used to illustrate the influence of NNIs is the conformational entropy. We used the obtained Gaussian model distributions in eq. 8 to calculate the conformational backbone entropy for GRG as well as for R1R2 in GRRG and for R1R2 and R2R3 pairs in GRRRG. The observed entropy values are listed in Table S3. Since the calculated entropy depends on the mesh size chosen for the probability density distribution (2° for both dihedral angles in our case), a reference system is needed.^33^ We use the 77.7 J/mol*K value which we calculated for an artificially constructed Ramachandran distribution of random coil type conformation which mimics the situation in which a residue samples the conformationally permissible space.^32^ As one can infer from the values in Table 2, the entropy values of all arginine based peptides investigated are considerably lower. For GRG, the obtained value of 67.45 J/mol*K corresponds to Helmholtz energy difference of ca. 3 kJ/mol. We observed a reduction of the entropy by -1.68 with regard to GRG and -0.65 J mol^-1^ K^-1^ for R1 and R2 of GRRG, which corresponds to a Helmholtz energy loss of only 492 and 190 J mol^-1^ at room temperature. For the pentapeptide, the entropy losses are -3.81, -3.2 and -3.46 J*mol^-1^ *K^-1^, which corresponds to a Helmholtz energy increase of 1.117, 0.938 and 1.014 kJ*mol^-1^ at room temperature. Compared with the above ‘random coil’ value, the corresponding entropy values indicate a reduction of the Helmholtz energy at room temperature by more than 4 kJ/mol. We can conclude that the NNI induced entropy reduction of arginine residues is significant in the pentapeptide but rather moderate in the tetrapeptide.

### MD simulations of GRRG and GRRRG

We wondered whether the obtained NNIs could be at least qualitatively reproduced by MD simulations. Identifying combinations of peptide/protein force fields and water models that can reproduce particularly experimental J-coupling constants has been attempted by several groups over the last ten years. Many of the canonical force fields have been optimized to account for the exceptionally high pPII propensity of alanine.^47,57^

**Figure 3.**
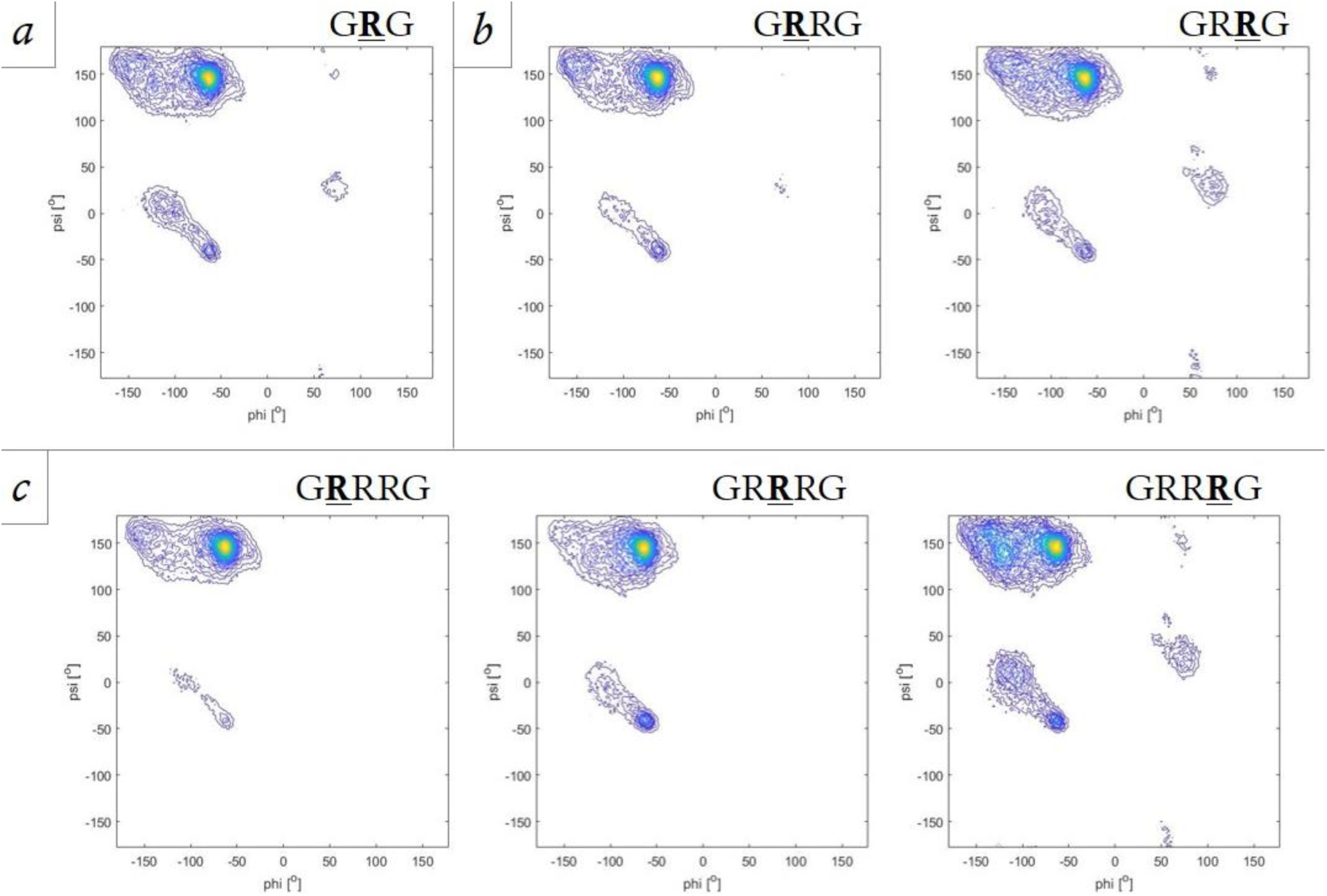
Ramachandran of arginine residues in GRG, GRRG and GRRRG obtained from MD simulations with an CHARMM36m force field combined with a TIP3P water model. Details of the simulations are described in Material and Methods.

Recently, Zhang et al. showed that Amber ff14SB combined with TIP3P reproduces experimental J-coupling constants and amide I’ profiles better than other Amber, OPLS and CHARMM force fields for guest alanine in GAG.^24^ Here, we first used Amber ff14SB and CHARMM36m with TIP3P and OPLS-AA/M with TIP4P to produce the Ramachandran distribution of cationic GRG. To access the obtained distributions, we calculated the respective reduced χ^2^-values for J-coupling constants and the VCD profile of amide I’ which are listed in Table S4. Apparently, CHARMM36m produces by far the best agreement with the experimental data. We therefore used this force field and TIP3P to produce conformational distributions for cationic GRG, GRRG and GRRRG. The corresponding Ramachandran plots are shown in Figure 3 and the mesostate occupations as a function of simulation time is shown in Figure S3. Figure S3 demonstrates the mesostate occupations remain relatively stable throughout the MD simulations. In order to compare them with the distributions obtained with our Gaussian models we calculated the occupation of the mesostates defined in Material and Methods. The mesostate occupations are listed in Table 3. As already indicated by the respective χ^2^-values (Table S4), the simulations capture the pPII and the overall β-strand propensities of GRG well. Contrary to the respective Gaussian distribution, however, the β-strand population is spread over a broader range of φ values thus covering β_t_, β_a_ and even β_p_. The respective occupations of the mesostate associated with right-handed helical/type III β turn structure are comparable. The comparatively small differences between MD and Gaussian distributions are also reflected in the differences between calculated and experimental J-coupling constant (Tables 1 and S4). The χ_J_^2^-value of 2.65 obtained for the MD distribution is larger than the Gaussian value, but the performance is very much comparable with what we observed for GAG.

**Table 3.** Mesostate populations of arginine residues in the Ramachandran plots of the indicated peptides as obtained from MD simulations with an CHARMM36m force field and a TIP3P water model.

|  |  | GRG | GRRG |  | GRRRG |  |  |
| --- | --- | --- | --- | --- | --- | --- | --- |
|  |  | *Ri* | *R1* | *R2* | *R1* | *R2* | *R3* |
| *pPll* | *Gaussian* |  | 0.46 | 0.3 | 0.42 | 0.21 | 0.3 |
|  | *MD* | 0.46 | 0.53 | 0.46 | 0.59 | 0.55 | 0.35 |
| $\beta_t$ | *Gaussian* | | 0.13 | 0.11 | 0.16 | 0.09 | 0.21 |
|  | *MD* | 0.11 | 0.11 | 0.14 | 0.11 | 0.13 | 0.19 |
| $a\beta$ | *Gaussian* | | 0.09 | 0.25 | 0.14 | 0.34 | 0.21 |
|  | *MD* | 0.12 | 0.13 | 0.11 | 0.12 | 0.06 | 0.12 |
| $p\beta$ | *Gaussian* | | 0.01 | 0.01 | 0 | 0 | 0.02 |
|  | *MD* | 0.06 | 0.05 | 0.07 | 0.05 | 0.07 | 0.04 |
| *rh* | *Gaussian* |  | 0.08 | 0.04 | 0.13 | 0.19 | 0.14 |
|  | *MD* | 0.05 | 0.05 | 0.03 | 0.02 | 0.07 | 0.04 |

An inspection of the MD Ramachandran plots of the arginine residues in GRRG and GRRRG (Figure 3) already indicates that the respective differences between the conformational distributions are much less pronounced than between the respective Gaussian distributions. Only for R3 of GRRRG the Ramachandran plot seems to indicate an enhanced sampling of β-strand conformations. This impression is confirmed by an inspection of mesostate populations. For GRRG, the total β-strand populations are 0.29 and 0.32. Corresponding values for GRRRG are 0.28, 0.26 and 0.43, indicating a significant increase of β-strand solely for R3. Not surprisingly, CHARMM36m produces much larger discrepancies between calculated and experimental J-coupling values, as indicated by the rather large χ_J_^2^-values (Table 1). Therefore, we must conclude that CHARMM36m, in spite of its capability to reproduce the experimental values of GRG comparatively well, significantly underestimates NNIs. This notion is further corroborated by Hellinger distances comparing Ri in GRG with the arginines in GRRG and GRRRG. They all lie between 0.11 and 0.18 indicting that the respective distributions are moderately similar. Our results seem to suggest that the electrostatic interactions between positively charged arginine groups do not seem to contribute substantially to the NNIs, since they are fully accounted for in the MD simulations.

### Estimating the length of poly-arginine peptides

The results of our Gaussian model-based analysis suggest that NNIs produce two effects. First it eliminates the populations of the conformations situated in the right half of the Ramachandran plot. This alone reduces the probability that a longer arginine chain would form turns and loops that are a necessity for compact structures. Second it causes a redistribution from pPII to β-strand and a shift of the latter towards more negative φ-values. If this occurs in poly-R segments as well it will lead to an increase of the end to end distances. Overall, the propensities of GRRRG listed in Table 1 seem to suggest an anticorrelation between the β-strand and the pPII population of nearest neighbors which Schweitzer-Stenner and Toal earlier reported for several GxyG tetrapeptides.^58^ We wondered to what extent poly-arginine with the arginine distribution of R2 in GRRRG would exhibit a 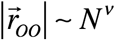 dependence where *N* is the number of residues and the exponent *ν* is indicative of the compactness/extension of the considered polymer. We calculated the *r*_*oo*_ as a function of the number of residues between the peptide groups using the formalism described in Material and Methods (eq. 12 and explanatory text). To this end we used the truncated distribution employed for the modelling of Raman amide I’ profiles. As shown in Figure 6, *r*_*oo*_ follows a power law with ν = 0.66 ± 0.02. The obtained statistical uncertainty suggests that the deviation from the 0.59 value for a self-avoiding random coil is significant. We performed the same calculation for poly-R segments assuming the conformational distribution of arginine in GRG. The corresponding plot is shown in Figure S4. The corresponding exponent if 0.62 ± 0.05. Thus, the margin of error includes the canonical 0.59 value and the 0.66 value obtained with our NNI-based calculation. Hence, we can conclude that while our NNI-based model clearly predicts a scaling behavior for poly-arginine which lies above the expectation for a self-avoiding random coil it is likely that this reflects a combination of intrinsic propensities of R and the influence of NNI. Mao et al.^16^ showed a scaling-law like behavior for plots of residue distances (averaged over all atoms and all peptide segments, i.e. all pairs, triplets, etc.) versus chain separation which is a measure of the number of considered residues, for a series of protamine sequences and poly-arginine. They did not report the respective ν-values. We simulated their curves with Matlab and found that the scaling exponent for their poly-arginine calculation is close to 0.7. This is not too distinct from the above prediction based on our conformational analysis of GRRRG.

**Figure 6:**
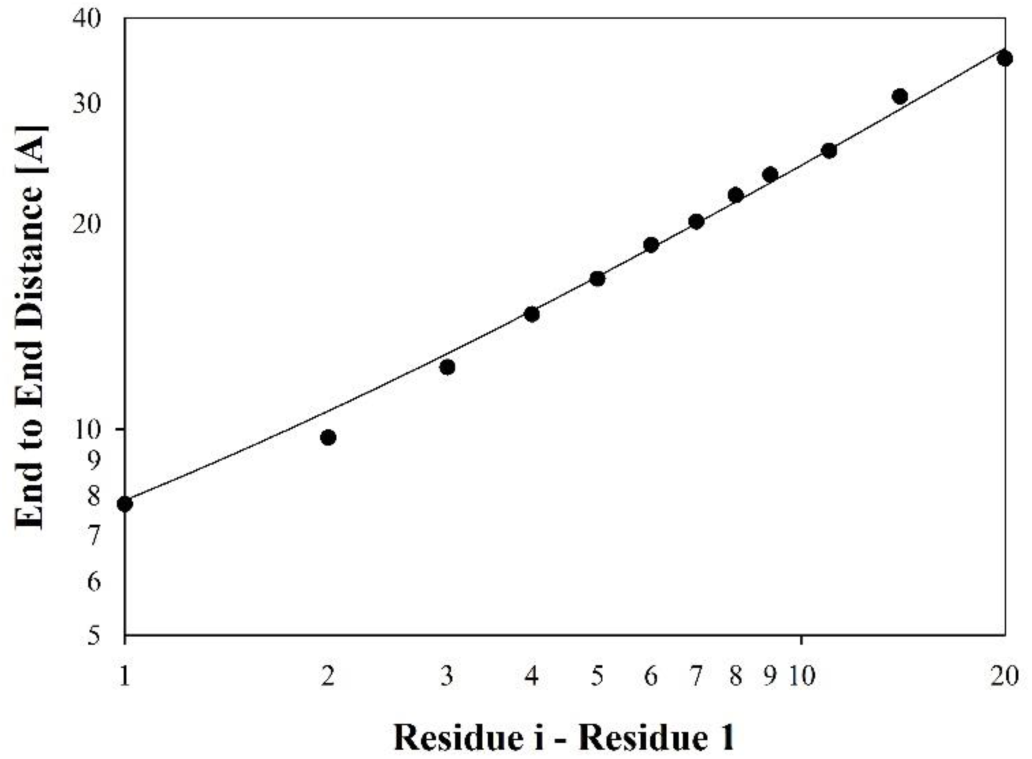
End to end distance between the N-terminal and the C-terminal carbonyl oxygen calculated as function of the difference between the number of residues of a hypothetical poly-arginine peptide for which we assumed the Ramachandran distribution obtained for the central residue R2 of GRRRG. The solid line results from a fit of a lower law to the calculated data.

Mao et al. related the extended structure of poly-arginine to electrostatic interactions. Their calculations clearly showed that the slope of the above described plots increase beyond the random coil level with increasing net charge of the considered peptide segment. However, in our case electrostatic interactions are unlikely to be relevant. At the chosen acidic pH value the chloride concentration is ca. 10 mM. If we take this as the ionic strength value the Debye length would be close to 0.64 nm. The real ionic strength is higher because of their have to be 3(4) counterions per peptide in solution. The total value for the ionic strength therefore varied between ca. 280 mM (for the Raman experiments with GRRRG) and 25 mM (for the NMR experiments with GRRG). Hence, the Debye length will be significantly shorter. As shown in Figure S5, representative distances between charged groups in the investigated peptides are above 1 nm, well outside of the range of electrostatic interactions.

We would like to emphasize at this point that our findings do not invalidate the role of electrostatics for long polypeptides or even proteins. It is obvious that electrostatic repulsion adds to the excluded volume effect and destabilizes compact conformations formed by non-local interactions. However, as demonstrated by the above calculation end to end distances depend on individual conformational distributions of residues which are determined by intrinsic propensities and local nearest neighbor interactions. Mao et al. reported a high probability that arginine samples the pPII conformation during MD simulations.^16^ For the poly-arginine case they report a probability of 0.23 for three and more arginine residue in a row to adopt a pPII conformation. This suggests an average pPII propensity of at least 0.61 for each residue. This value is very close to the pPII propensities that emerged from our analysis of GRG and of the R1 residues of GRRG and GRRRG. (Table 1), but significantly larger than respective values obtained for R2 and R3 of these peptides.

The corresponding counting of β-strand conformations starts with two consecutive residues. Thus, the 0.08 value reported for poly-arginine correspond to a minimal propensity of 0.28. This value is higher than the 0.2 obtained from our Gaussian analysis of GRG, but very close to the values obtained for the R1 residues of GRRG and GRRRG, but significantly lower than the β-strand propensities of R2 and R3 in both peptides. For right-handed helical conformations Mao et al. reported that propensity values for helical trimers are negligible low. This is not at odds with our own per residue propensities. Even the obtained values for GRRRG would yield an average value of 0.006 for three helical conformations to exist in a row. Overall, we can conclude that the MD-simulations of Mao et al. might overestimate the pPII content of polyarginines similarly to what we observed with our own MD simulations (*vide supra*). However, it deserves to be noticed in this context that the pPII trough considered by Mao et al. is larger than ours (−120° < ϕ < -60°). Hence, it covers part of the βt mesostate which we count towards the β-strand sampling.

## CONCLUSION

The Gaussian model analysis of J-coupling constants and amide I’ Raman band profiles of GRRG and GRRRG revealed a rather strong nearest neighbor interaction between arginine residues which eliminates conformations in the right half of the Ramachandran plot and redistributes conformational sampling from pPII to β-strand. Additionally, they shift the β-strand basin to more negative values. Thus, they cause a more extended average structure of the respective peptides. A Hellinger distance-based analysis reveals that the R-residues in both peptides differ more from that in GRG and they do among each other. However, a calculation of the conformational entropy shows that a more significant reduction in GRRRG due to NNIs. A comparison of individual propensities in the latter suggest an anticorrelation between pPII and β-strand propensities, in line with earlier observations.^58^ In view of the ionic strength values of our samples it is unlikely that the observed NNIs are electrostatic in nature. They are most likely solvent mediated as predicted by Avbelj et al.^26,59^ This notion is corroborated by much less pronounced NNI effects obtained with MD simulations using CHARMM36m + TIP3P. These calculations should fully account for any electrostatic effects but are likely to underestimate strongly cooperative interactions between solvent molecules which give rise to solvent-mediated NNIs. We used the conformational distribution of R2 in GRRRG to predict the length of the polyarginine as a function of the number of residues and found that it follows a power law with a scaling coefficient of 0.66, which is statistically significantly larger than the 0.59 value expected for an ideal random coil with excluded volume. We argue that our results show the relevance of individual propensities and their dependence on NNIs for the conformational sampling of peptides and proteins in aqueous solutions which do not adopt collapsed states (ν>0.5). In line with an earlier suggestion of Scheraga and coworkers we suggest to use the term statistical coil for the conformational manifolds of such systems.^60^

## Associated content

The following tables and figures can be found in the Supporting Material:, Karplus plots for the utilized coupling parameters, a visualization of mesostates, a plot of the end to end distance between carbonyl oxygens of a hypothetical poly-arginine peptide, visualizations of inter-residue distance in extended GRRG and GRRRG and a table listing conformational entropies.

## Author contributions

B.M has performed the reported NMR and Raman experiments. B.A. has performed the MD simulations. B.M. and R.S.S. have analyzed the experimental data and wrote most of the paper. B.A. wrote part of the MD section of the paper. H.S and B.U. have supervised the NMR experiments and MD simulations and contributed to the discussion of the data

## Acknowledgements

This project was supported by grant from the National Science Foundation to B.U. and R.S.S (MCB-1817650). B.M. acknowledges a travel grant from the International Office of Drexel University to conduct research in Frankfurt, Germany. Work at BMRZ is supported by the state of Hesse. The writing of the paper required some private support from R.S.S. for office equipment which could not be purchased by institutional order because of the pandemic.

